# Comparative Transcriptomics Reveals Context- and Strain-Specific Regulatory Programs of *Agrobacterium* During Plant Colonization

**DOI:** 10.1101/2025.05.27.656240

**Authors:** Yu Wu, Hsin-Yi Chang, Chih-Hang Wu, Erh-Min Lai, Chih-Horng Kuo

## Abstract

*Agrobacterium* is a genus of plant-associated bacteria capable of transferring DNA into host genomes to induce tumorigenesis. The process has been primarily studied in a few model strains, particularly C58, and developed into *Agrobacterium*-mediated transformation (AMT) for genetic manipulation. However, the diversity of wild-type strains and their context-specific regulatory responses remain poorly characterized. Here, we evaluated five wild-type strains and identified 1D1108 as superior in tumorigenesis on legumes and transient transformation in *Nicotiana benthamiana*. Under *in vitro* virulence induction with acetosyringone (AS), we identified 126 differentially expressed genes (DEGs) in 1D1108. Although the number of DEGs was comparable to those in C58 and the legume isolate 1D1609 under the same condition, only 22 DEGs, primarily within the *vir* regulon, were conserved, indicating extensive divergence among these *Agrobacterium* strains. Leaf infiltration of *N. benthamiana* revealed 1,134 DEGs specifically regulated *in planta* for 1D1108. These included genes involved in attachment, virulence regulation, type IV pilus, succinoglycan biosynthesis, and diverse nutrient transporters, providing new evidence on expression regulation during colonization. Comparative analyses of *in planta* transcriptomes with C58 and *Pseudomonas syringae* DC3000 revealed distinct secretion systems required for pathogenesis, namely type IV for *Agrobacterium* and type III for *Pseudomonas*, and only approximately 5-19% of DEGs were conserved. These limited transcriptomic overlaps underscore the importance of studying gene expression in strains and conditions directly relevant to the biological context, rather than relying on model systems. Together, this work reveals how environmental and host-associated cues shape transcriptional responses in plant-associated bacteria.

**IMPORTANCE:** *Agrobacterium* is a key tool for plant genetic engineering, yet much of our knowledge about its biology comes from a few strains studied under artificial conditions. This work combines tumor formation assays on legumes and transient transformation in *Nicotiana benthamiana* with transcriptomic analyses to explore how diverse *Agrobacterium* strains, particularly the high-performing wild-type strain 1D1108, respond to host and environmental cues. By comparing gene expression under *in vitro* and *in planta* conditions, we found that the bacterial response within plant tissue is more complex than previously appreciated. Cross-strain comparisons further revealed limited conservation in gene expression regulation, even under similar conditions. Comparative analysis with *Pseudomonas syringae* revealed the activation of distinct secretion systems for pathogenesis and differences in regulatory programs used during plant colonization. These findings underscore the importance of studying bacteria in biologically relevant contexts and highlight the limitations of relying solely on model strains to infer regulatory responses.

## INTRODUCTION

The genus *Agrobacterium* contains a group of soil bacteria associated with plants and are notable for their capacity of causing crown gall or hairy root diseases (Nester, 2015; de Lajudie et al., 2019). The phytopathogenicity of these bacteria depends on oncogenic plasmids that are either tumor-inducing (pTi) or root-inducing (pRi) (Weisberg et al., 2020, 2022). During the infection process, a specific segment of transfer DNA (T-DNA) located on the oncogenic plasmid is processed and transported into the host cells via the type IV secretion system (T4SS) encoded on the same plasmid (Hwang et al., 2017; Weisberg et al., 2023). The T-DNA can be integrated into the plant nuclear genome, leading to expression of T-DNA genes for the biosynthesis of plant hormones and opines. The transformation not only results in abnormal proliferation of plant cells, but also converts the plant cells into food factories for the invading bacteria.

This unique interkingdom DNA transfer has been exploited by biologists to develop *Agrobacterium*-mediated transformation (AMT) as a powerful tool for genetic manipulation of plants (Kado, 2014; Hwang et al., 2017). Compared to the gene gun method, which uses microparticle bombardment, AMT has several key advantages including low cost, easy to use, and minimal damages to the target genome. Moreover, in addition to stable transformation, AMT can also be used for transient expression of genes encoded by T-DNA without integration. Therefore, AMT offers flexibility for the study of gene functions in plants and the generation of genetically modified organisms for biotechnology applications.

To develop AMT for genetic engineering, many studies have been devoted to investigate the molecular mechanisms and genes involved (Hwang et al., 2017; Weisberg et al., 2023). The infection process starts from the attachment of bacterial cells to the host, which involves beta-1,2 glucan production by ChvB and transport by ChvA (Zorreguieta et al., 1988; Cangelosi et al., 1989). Effective infection involves recognition of plant wounding signals, including acidity at the pH range of 5.5-6.0 commonly found in plant apoplast (Geilfus, 2017), as well as monosaccharides and phenolic compounds such as acetosyringone (AS) (Stachel et al., 1985). These signals can be sensed by bacterial two-component systems, such those encoded by *chvG*/*chvI* for pH change (Li et al., 2002) and *virA*/*virG* for phenolics (Lee et al., 1995), leading to the expression of virulence (*vir*) genes located on oncogenic plasmids. Subsequently, T-DNA is processed and transferred into the host cells by various Vir proteins, including VirB1-11 and VirD4 that together constitute a functional T4SS (Lai and Kado, 2000).

Despite the advancements, two critical knowledge gaps remained regarding agrobacterial gene expression regulation. First, the genus *Agrobacterium* harbors extensive biological diversity, but studies on virulence have focused on only a few strains. To date, more than 20 genomospecies have been defined based on genomic divergence (Weisberg et al., 2023). Strains belonging to the same genomospecies may differ by up to 15% of their gene content, while strains belonging to different genomospecies may differ by more than 20% in gene content (Haryono et al., 2019; Wu et al., 2019; Chou et al., 2022). In addition to the diversity of chromosomal genes, pTi can be classified into at least 11 types (Weisberg et al., 2020, 2022; Chou et al., 2022). Importantly, the genetic diversity is likely linked to phenotypic diversity such as host range and transformation efficiencies (Hwang et al., 2013). To date, most studies on agrobacterial virulence have focused on strain C58 belonging to genomospecies 8, which is also the wild-type progenitor of nearly all disarmed strains commonly used for AMT such as GV3101 and EHA105 (Kado, 2014; Hwang et al., 2017). Second, for transcriptomics investigation, most previous studies not only focused on C58, but also mainly examined *in vitro* conditions (Yuan et al., 2008; Wilms et al., 2012; Lee et al., 2013; Heckel et al., 2014; Möller et al., 2014; Dequivre et al., 2015; Raja et al., 2018). Two exceptions include one that compared C58 to a genomospecies 7 strain 1D1609 for their responses to *in vitro* AS treatment by RNA-Seq (Haryono et al., 2019) and one that investigated C58 gene expression inside *Arabidopsis thaliana* tumors by microarrays (González-Mula et al., 2018).

To bridge the aforementioned gaps and to expand our understanding of *Agrobacterium* biology, we conducted tumorigenesis and transformation assays of five wild-type *Agrobacterium* strains (**Table 1**). These strains differ from C58 in their host range according to artificial inoculation experiments (Hwang et al., 2013). Also, our genomic characterization revealed their divergence in genomospecies assignment, gene content, and pTi type (Wu et al., 2019; Chou et al., 2022). We discovered that strain 1D1108 consistently exhibited high efficiencies of inducing tumor formation in kidney bean and soybean, as well as a high transient transformation efficiency in *Nicotiana benthamiana*. Therefore, we conducted RNA-Seq experiments to investigate 1D1108’s transcriptomic responses to acidic environment and AS treatment under *in vitro* conditions, as well as under an *in planta* condition following infiltration into *N. benthamiana* leaves. Through these experiments, together with comparisons with previous *Agrobacterium* transcriptomics results (Yuan et al., 2008; González-Mula et al., 2018; Haryono et al., 2019) and *in planta* transcriptomics of *Pseudomonas syringae* (Lovelace et al., 2018; Nobori et al., 2018), we identified the commonalities and specificity in gene expression regulation among diverse plant pathogens in response to different environmental cues.

**Table 1:** The *Agrobacterium* strains used in this study. The genomospecies and pTi type assignment are based on Weisberg et al. 2020 (DOI: 10.1126/science.aba5256).

| Strain | Natural Host | NCBI Accession | Genomospecies | pTi Type | Opine Type | Reference |
| --- | --- | --- | --- | --- | --- | --- |
| 1D1108 | *Euonymus* | GCA_003666425.1 | G1 | I.b | Nopaline | Wroblewski et al. (2005); DOI:<br>10.1111/j.1467-7652.2005.00123.x |
| 1D1460 | Raspberry vine | GCA_003666445.1 | G4 | I.a | Nopaline | Wroblewski et al. (2005); DOI:<br>10.1111/j.1467-7652.2005.00123.x |
| 1D1609 | Alfalfa | GCA_002943835.1 | G7_a | II | Octopine | Palumbo et al. (1998); DOI:<br>10.1007/s002030050586 |
| 1D132 | Cherry | GCA_003667725.1 | G8_a | I.b | Nopaline | Wroblewski et al. (2005); DOI:<br>10.1111/j.1467-7652.2005.00123.x |
| Ach5 | Yarrow | GCA_000971565.1 | G1 | II | Octopine | Archdeacon et al. (2000); DOI:<br>10.1111/j.1574-6968.2000.tb09156.x |
| C58 | Cherry | GCA_000092025.1 | G8 | I.a | Nopaline | Lin and Kado (1977); DOI:<br>10.1139/m77-229 |

## RESULTS AND DISCUSSION

### *Agrobacterium* strain 1D1108 is highly efficient in stable and transient transformation

To evaluate tumorigenesis efficiencies of the selected *Agrobacterium* strains, we measured tumor formation rates and weights at five weeks post-inoculation. The five wild-type strains examined in this work (i.e., 1D1108, 1D1460, 1D1609, 1D132, and Ach5) differed significantly in their performance against two legume hosts (**Fig. 1**). For inoculation in kidney bean, 1D1108 and 1D1460 both showed > 90% tumor formation rates averaged across three batches, which are significantly higher than the other three strains (**Fig. 1B**). In soybean, 1D1108 and 1D1069 showed > 90% tumor formation rates, significantly higher than the other three strains (**Fig. 1E**). For tumor weight, 1D1108 was capable of inducing significantly larger tumors than all other four strains in both hosts (**Fig. 1C and 1F**). Our quantitative assays using both metrics revealed that 1D1108 was the best performing strain in both hosts. 1D1460 and 1D1609 also showed high virulence on both hosts, with 1D1609 as the second best in soybean and 1D1460 second to 1D1108 in kidney bean. Our assays also revealed the induced tumors on soybean were approximately one order of magnitude smaller. Nevertheless, those induced tumors were still clearly visible upon visual inspection.

**FIG. 1.**
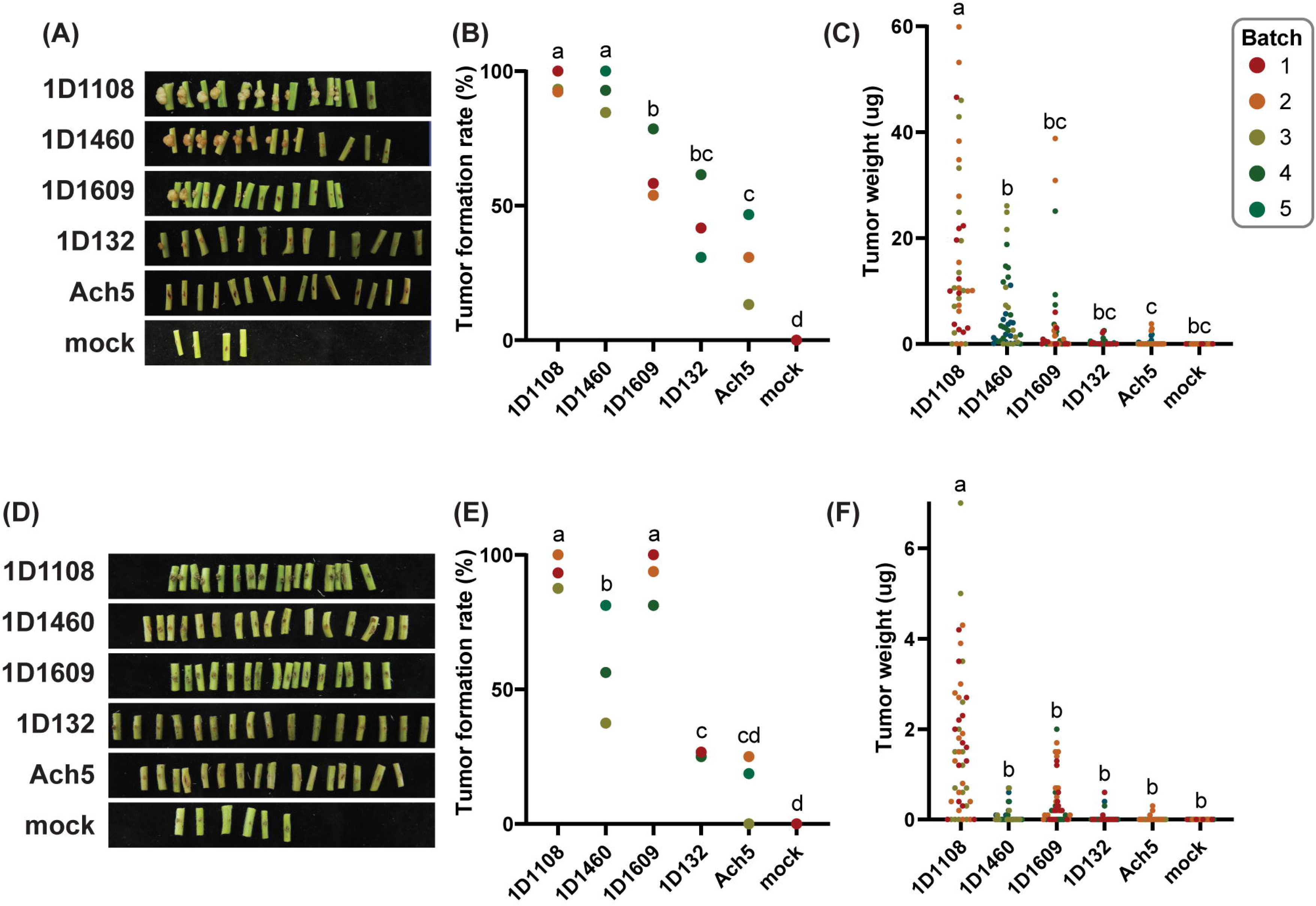
Tumor formation induced by wild-type *Agrobacterium* strains. (A) Representative photos, (B) tumor formation rates, and (C) tumor weight distribution on kidney bean. (D) Representative photos, (E) tumor formation rates, and (F) tumor weight distribution on soybean Tainan No. 7. The results were collected at five weeks after inoculation. A total of five batches, with sample sizes ranging between 11 to 16 plants per batch, were used. Each strain was tested in three different batches. The tumor formation rates among strains were compared using one-way ANOVA and Tukey post-hoc test. For tumor weight, data points that deviate from the mean by more than three standard deviations were considered as outliers and excluded.

For transient transformation based on agroinfiltration into mature leaves, we utilized the RUBY reporter system (He et al., 2020). The expression of this reporter converts tyrosine in plant cells to the red pigment betalain, enabling precise quantification by absorbance. Our preliminary tests using kidney bean and soybean produced results that were generally weak, uneven within the infiltrated regions, and variable among leaves. Therefore, *N. benthamiana* was selected for this assay. To control for variability among individual leaves and across batches, C58 was infiltrated in each leaf for normalization. Based on approximately 30 samples distributed across three batches for each strain, 1D1108 performed significantly better than the other four wild-type strains, averaging 3.8-fold higher signal than the reference strain C58 (**Fig. 2**).

**FIG. 2.**
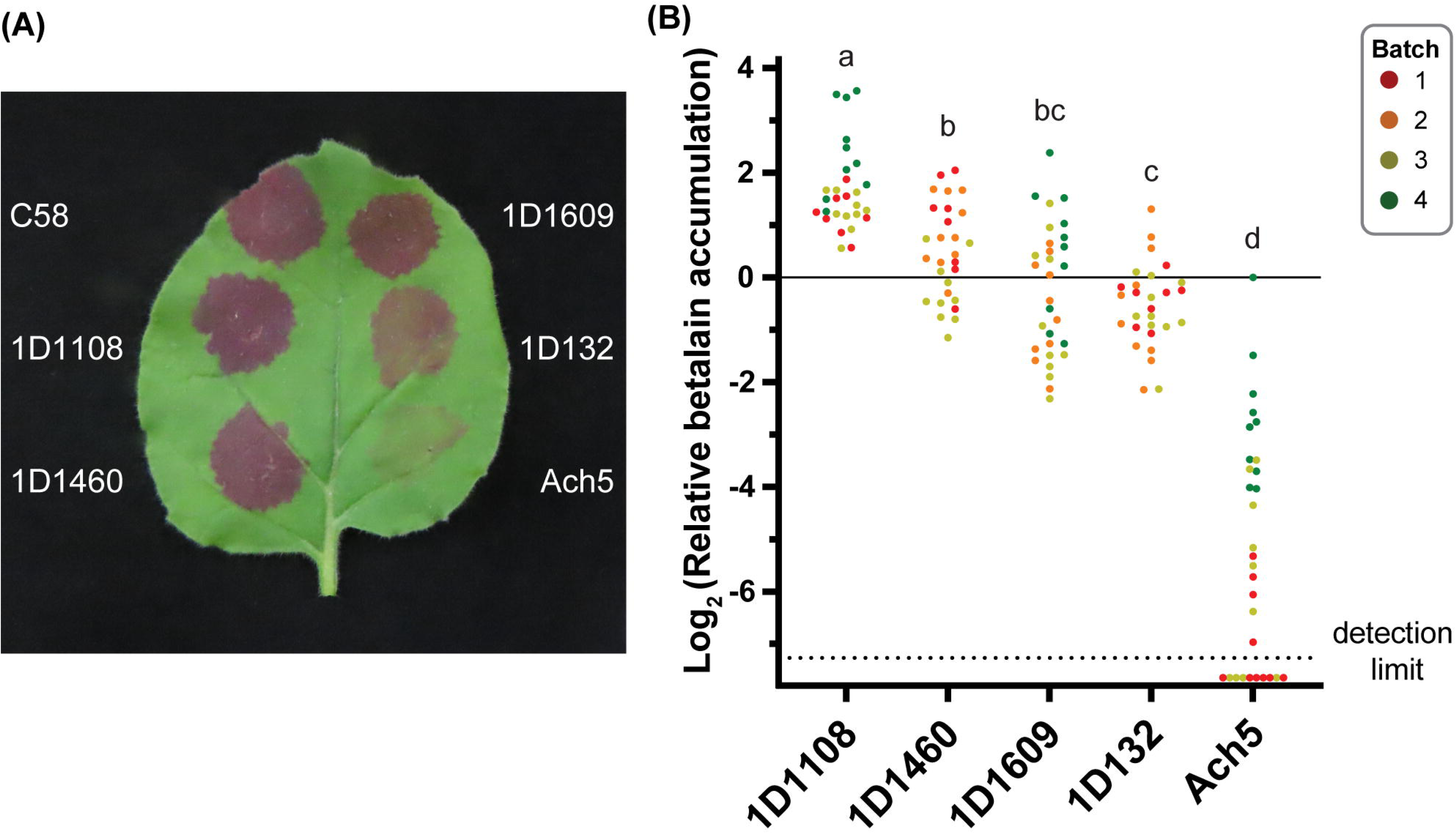
Transient transformation by wild-type *Agrobacterium* strains. (A) A representative photo and (B) relative betalain accumulation levels normalized to the reference strain C58. Leaves of 30-day old *Nicotiana benthamiana* were used for infiltration, results were collected at 48 hours post infiltration. A total of four batches, with sample sizes ranging between 8 to 10 leaves per batch, were used. Each wild-type strain was tested in three different batches. The reference strain C58 was infiltrated into each leaf and used for normalization. Data points with a raw betalain accumulation value of zero or below were plotted under the dotted line that indicates the detection limit. The relative betalain accumulation levels among strains were compared using one-way ANOVA and Tukey post-hoc test.

In summary, our results from stable and transient transformation assays demonstrate that 1D1108, a wild-type strain that has received limited prior study, has strong potential to reveal new insights into AMT and support future improvements.

### Implementation of *in vitro* and *in planta* RNA-Seq experiments

To investigate the genes contributing to the high efficiencies of 1D1108, we conducted RNA-Seq experiments to characterize its transcriptomic responses to conditions relevant to AMT. These included four conditions, each with three biological replicates (**Fig. 3A**). Condition A, axenic culture in the minimal medium at pH 7.0, was used as the baseline. Conditions B and C were used to investigate the effects of two key regulators of virulence-related signal transduction, namely an acidic environment (i.e., pH 5.5) and the presence of a phenolic compound (i.e., AS), respectively. Importantly, we established an *in planta Agrobacterium* transcriptome corresponding to the transient transformation in *N. benthamiana* in condition D. For this, we homogenized the infiltrated leaves at 16 hours post-inoculation (hpi), and enriched bacterial cells via filtration and centrifugation prior to RNA extraction.

**FIG. 3.**
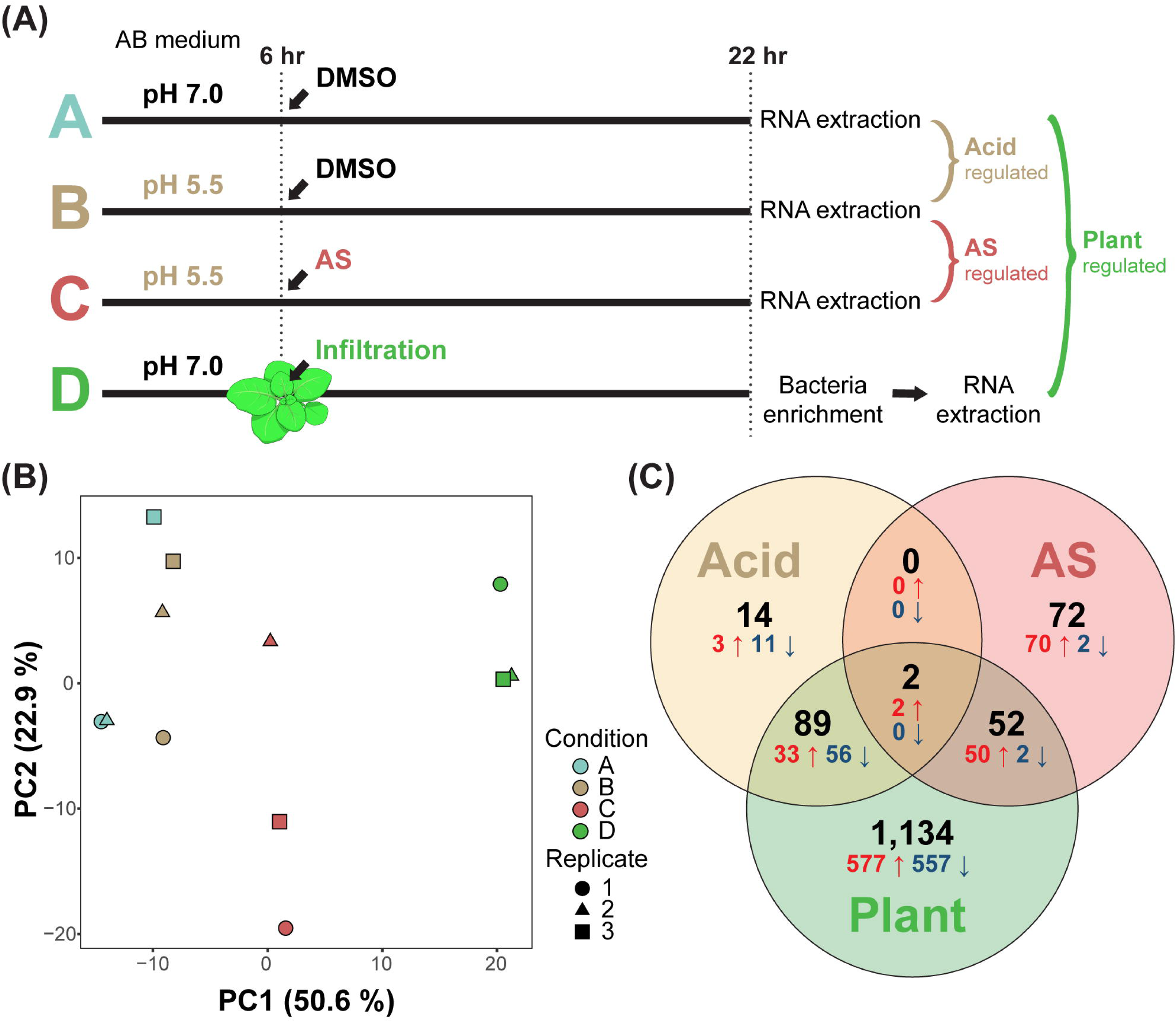
Design and summary results of the RNA-Seq experiment. (A) Conditions, time points, and pairwise comparisons of the RNA-Seq experiment. (B) Principal component analysis. Dots representing individual samples are color-coded according to the condition. Distance between dots indicates the level of dissimilarity. The percentage of variance explained by each axis is provided in parentheses. (C) A Venn diagram that illustrates the distribution of differentially expressed genes (DEGs; |log_2_(fold change)| ≥ 1 and *p*-adjusted < 0.01). Among the 5,359 protein-coding genes in strain 1D1108, a total of 1,353 were differentially expressed in at least one of the pairwise comparisons between conditions. The combined numbers of DEGs were indicated in black, while the numbers of up- and down-regulated genes were indicated in red and blue, respectively.

The 150-bp paired-end Illumina sequencing generated on average approximately 7.3 million quality-filtered reads per sample for the three *in vitro* conditions (i.e., A, B, and C) (**Table S1**). For condition D, on average, approximately 38.0 million quality-filtered reads were obtained per sample. Among these, approximately 15.2 million reads (40%) were mapped to the genome sequence of 1D1108, and the remaining reads were almost entirely mapped to the plant genome. These results demonstrated that our bacterial enrichment process in condition D was effective. Importantly, all 12 samples exceeded the recommended sequencing depth of 2 to 3 million reads for bacterial RNA-Seq (Haas et al., 2012).

Principal component analysis revealed that differences in gene expression across conditions were primarily explained by the first principal component (PC1), which accounted for 50.6% of the total variance (**Fig. 3B**). In contrast, variation among biological replicates was mostly captured by PC2, explaining 22.9% of the variance. These results indicate that environmental condition was the main driver of gene expression patterns. Notably, the *in planta* transcriptome was clearly distinct from all *in vitro* conditions, highlighting extensive transcriptional reprogramming in *Agrobacterium* when colonizing plants.

### *In planta* transcriptome reveals shared and unique differentially expressed genes

Among the 5,359 protein-coding genes in strain 1D1108, 1,353 (25.2%) were identified as differentially expressed genes (DEGs), defined by |log_2_(fold change)| ≥ 1 and *p*-adjusted < 0.01, in at least one of three pairwise comparisons (**Fig. 3C; Table S2**). The acidic environment (B vs A; “Acid”) resulted in 105 DEGs. Phenolic induction (C vs. B; “AS”) yielded a comparable number of DEGs, with 126 identified. In contrast, the plant environment (D vs. A; “Plant”) resulted in 1,277 DEGs.

In the plant environment, dozens of DEGs overlapped with those induced by either acidity or AS (**Fig. 3C; Table S2**). These included acid-induced genes encoding the type VI secretion system (T6SS) for interbacterial competition (Wu et al., 2012, 2019; Ma et al., 2014), as well as AS-induced *vir* and T4SS genes involved in T-DNA transfer (Haryono et al., 2019). Notably, *Agrobacterium* cells used for *in planta* transcriptome were resuspended in infiltration buffer without AS, mimicking natural infection condition. Thus, such co-induction suggests that machinery for both competition and transformation is activated in *Agrobacterium* during plant colonization. However, 1,134 DEGs were specific to the “Plant” set, underscoring the extensive regulatory responses to host-derived cues that are not elicited by acidity or AS alone.

To explore these responses further, we conducted functional enrichment analysis using the Clusters of Orthologous Genes (COG) classification (**Fig. 4; Table S2**). The most enriched categories among genes up-regulated inside the plant were “W” (extracellular structures, 37.5%; mainly type IV pili) and “U” (intracellular trafficking, secretion, and vesicular transport, 36.1%; mainly T4SS). By gene count, category “G” (carbohydrate transport and metabolism) was the most represented, with 65 out of 322 genes up-regulated *in planta*. These findings illustrate the breadth of transcriptional reprogramming triggered by plant colonization.

**FIG. 4.**
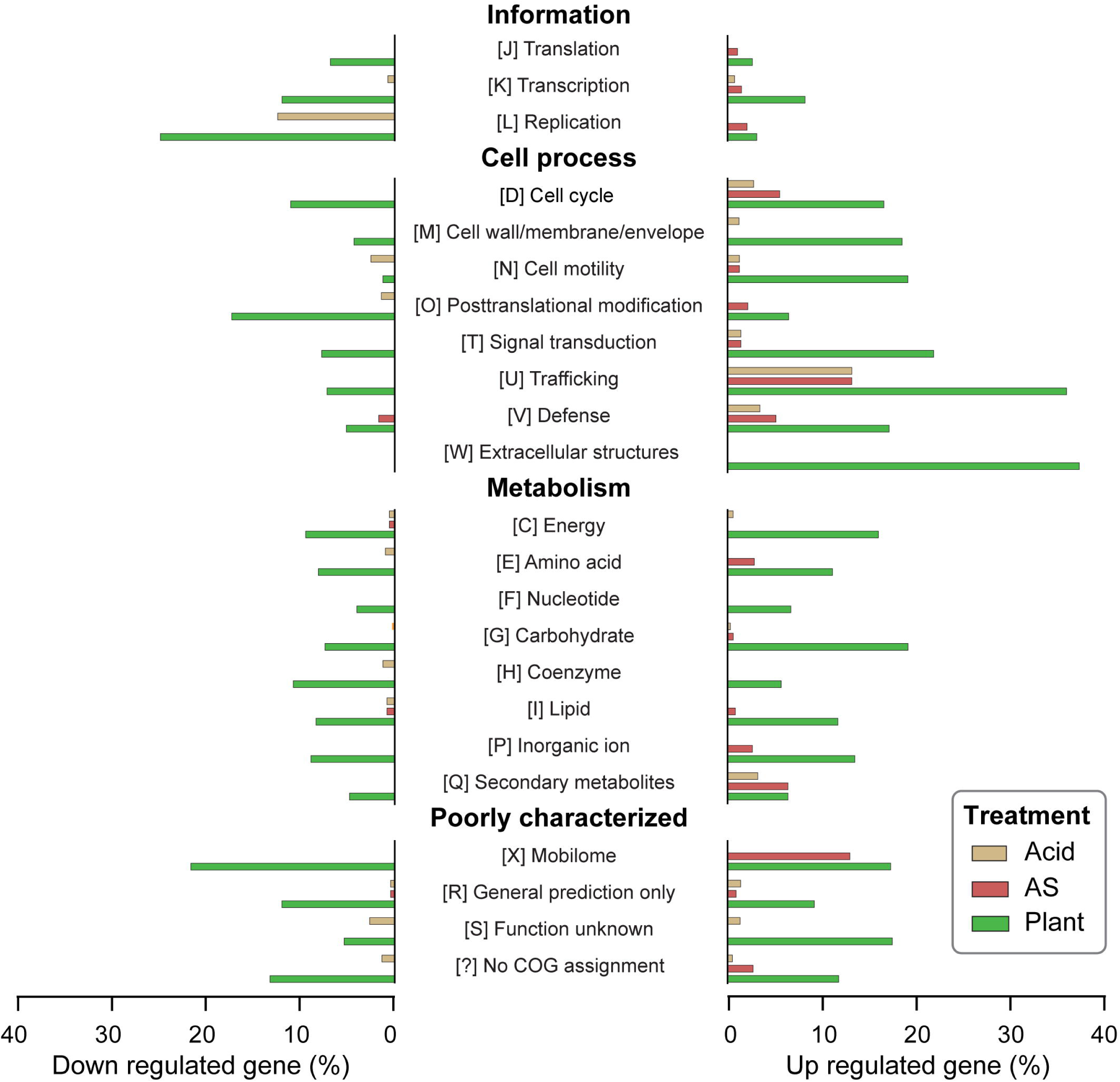
Functional category enrichment of differentially expressed genes (DEGs). The functional category assignments were based on the Clusters of Orthologous Genes (COG). The percentages of genes that were up- and down-regulated in each functional category were plotted for the three treatments.

### Distinct transcriptomic profiles among replicons

To assess how differential gene expression is organized at the genome level, we examined the distribution of DEGs across the replicons of 1D1108 and identified distinct patterns under the three treatments (**Fig. 5A**). Acidity did not result in preferential induction of DEGs across the replicons, apart from the T6SS gene cluster located near one end of the linear chromid (**Fig. 5B**). In contrast, AS induction led to a focused up-regulation of genes on pTi, particularly within the *vir* regulon. The plant environment triggered a more complex response: up- and down-regulated genes were observed in similar proportions on both the chromosome and chromid, while the accessory plasmids pAta and pAtb showed predominantly down-regulation. Notably, pTi exhibited strong induction under both AS induction and the plant environment, consistent with its role in virulence (**Fig. 5C**).

**FIG. 5.**
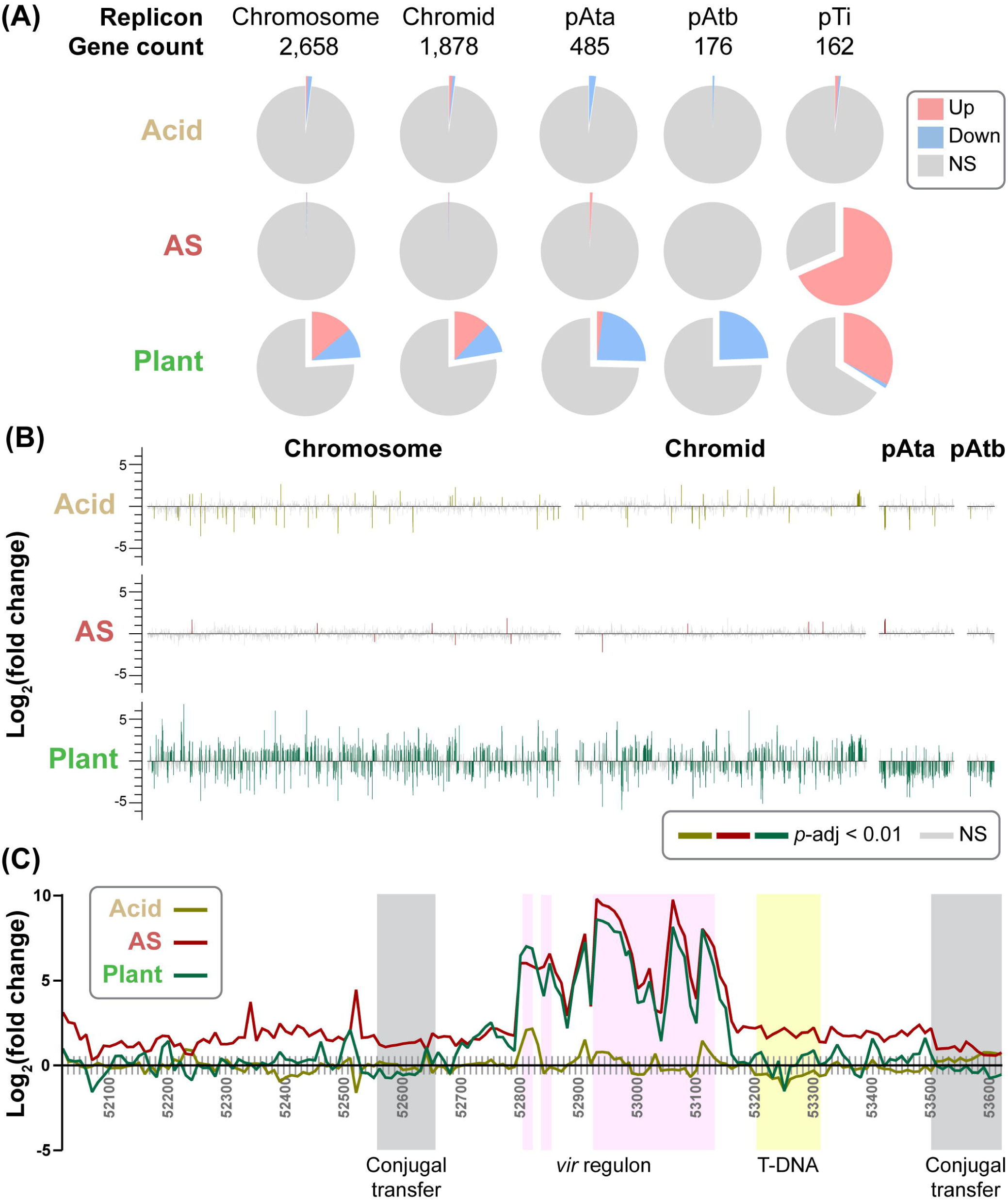
Analysis of the 1D1108 transcriptome based on genomic locations. (A) Distribution of differentially expressed genes (DEGs; |log_2_(fold change)| ≥ 1 and *p*-adjusted < 0.01) among replicons under different environmental cues. The pie charts illustrate the proportion of genes that are up- or down-regulated in each replicon under the specified factor. Abbreviation: NS, not significant. (B) and (C): relative positions and expression levels of genes on each replicon. Within each replicon, all protein-coding genes were plotted in the same size and arranged based on their relative positions, non-coding regions were omitted. For panel (C), genes with known functional significance were shaded with background colors for highlighting (conjugal transfer: gray; *vir*: pink; T-DNA: yellow).

A closer examination of pTi revealed over 20 genes in the *vir* regulon were up-regulated in response to AS induction and the plant environment (**Figs. 5C and 6; Table S2**). These included regulatory genes *virA* and *virG*, along with structural and processing components such as *virB1–11*, *virD1–5*, and *virE1–3*, mostly essential for T-DNA mobilization (Weisberg et al., 2023). Additionally, the *accABCDEFG* operon located immediately upstream of the *vir* regulon and involved in opine transport (Kim and Farrand, 1997) was up-regulated under acidic and plant environment.

**FIG. 6.**
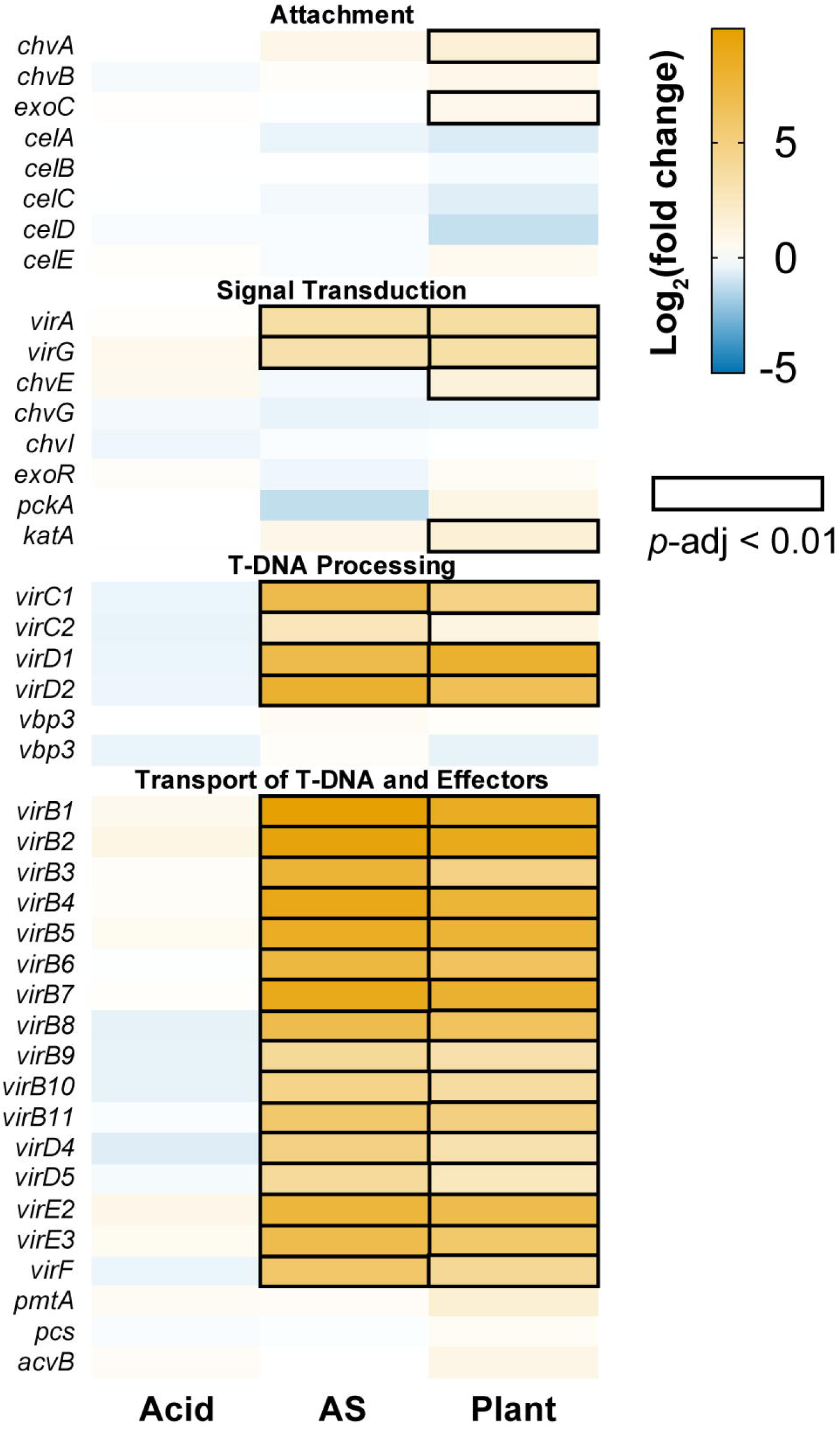
Differential expression of genes related to *Agrobacterium* virulence. The relative expression levels were plotted as a heatmap. Genes that were up- or down-regulated were plotted in orange or blue, respectively. The differential expression levels that reached statistical significance (*p*-adjusted < 0.01) were indicated by black boxes.

Interestingly, only two genes, *virH1* and *virH2* encoding P450-type monooxygenases, were consistently up-regulated under all three conditions (**Fig. 3C**). These two genes are not essential for virulence but are likely involved in the detoxification of antimicrobial phenolic compounds, including the *vir* inducers (Kalogeraki et al., 1999).

Unexpectedly, many genes located in other regions of pTi, including those within the T-DNA, were induced by AS but not by the plant environment. Unexpectedly, many genes located on other pTi regions, including those in T-DNA, were induced by AS but not the plant environment. This discrepancy may reflect prior observations that AS can increase pTi copy number through elevated RepABC protein expression (Cho and Winans, 2005). In contrast, phenolic signals may be less abundant in the apoplast of *N. benthamiana*, or other plant-derived factors may suppress pTi replication.

While AS and plant colonization both activated *vir* genes on pTi, several relevant chromosomal genes, such as the *celABC* and *celDE* operons for cellulose fibril synthesis, did not respond to either stimulus (**Fig. 6**). These genes are not essential for virulence, but can promote virulence by anchoring agrobacteria to plant cells (Matthysse et al., 2005). Additionally, the *chvG*/*chvI* two-component system and its periplasmic repressor *exoR*, which sense acidity (Wu et al., 2012), showed constitutive expression across all conditions.

### Known and novel virulence-associated genes regulated *in planta*

In addition to the *vir* regulon genes on pTi, which were described in the previous section, we identified other known virulence-associated genes that were specifically up-regulated during plant colonization. These results further support the biological relevance of the *in planta* transcriptome. At the same time, we discovered a broad set of genes not previously associated with virulence regulation, offering new insights into *Agrobacterium*-host interaction.

Among the known factors, we observed up-regulation of *chvA* and *exoC*, chromosomal genes involved in the synthesis and export of β-1,2-glucan required for bacterial attachment to plant (Zorreguieta et al., 1988; Cangelosi et al., 1989; Uttaro et al., 1990) (**Fig. 6**). Also up-regulated were *chvE*, encoding a sugar-binding protein that modulates the VirA/VirG signal transduction system (Doty et al., 1993; Shimoda et al., 1993), and *katA*, encoding a catalase that detoxifies reactive oxygen species and enhances virulence (Xu and Pan, 2000; Xu et al., 2001).

Beyond these well-characterized genes, we identified other genes previously unknown to be regulated by plant signals and lack evidence for their role in virulence. These include *ctpABCEF*, encoding a type IV pilus involved in reversible surface attachment (Wang et al., 2014); *exoACHKMNOPQVY* for succinoglycan biosynthesis (Matthysse, 2018); and *cspA*, a cold shock protein recognized as a microbe-associated molecular pattern in age-dependent plant immunity (Saur et al., 2016). Several transport systems were also induced, including those for alpha-glucoside (*aglAEFGK*), fructose (*frcABC*), glycerol (*glpPQSTV*), phosphonate (*phnCDEH*), phosphate (*pstABCS*), and sn-glycerol-3-phosphate (*ugpABE*).

Genes down-regulated within the plant environment included those involved in cell division (*ftsEHI* and *mraZ*), thiamine biosynthesis (*thiCGO*), conjugal transfer (*traABCDFGH* and *trbBKL*), and the transport systems for alpha-1,4-digalacturonate (*aguEFG*), molybdate (*modAC*), ribose (*rbsABC*), and zinc (*znuAC*).

Together, these expression patterns demonstrate that, beyond the activation of known virulence genes, plant colonization involves broader transcriptional reprogramming. These findings reveal candidate genes potentially involved in early stages of infection and provide a foundation for future functional studies.

### Shared virulence and strain-specific gene expression in *Agrobacterium*

To investigate the commonalities and differences in gene expression regulation among diverse *Agrobacterium* strains, we compared our RNA-Seq results to three published transcriptomics studies. First, we focused on the response to acidity. Since our *in vitro* pH 5.5 treatment was designed to mimic the plant apoplast, we included the “Plant” dataset for comparison. The reference study was a microarray-based investigation of C58 grown at pH 5.5 in axenic culture (Yuan et al., 2008). Strains 1D1108 and C58 belong to different genomospecies in the same genus and have similar genome sizes, each encoding about 5,350 protein-coding genes. We identified 4,173 single-copy homologs conserved between these two strains (**Table S3**). Among these genes, only 10 showed consistent differential expression in all three datasets (**Fig. 7A**), including one sulfatase, two *vir* (*virH1/H2*) and seven T6SS genes (*impBCDFGIJ*) (**Table S3**). This comparison revealed substantial differences in acid-responsive regulation between the strains. For example, the acidic environments induced differential expression of 72 genes in 1D1108 but not C58, these include 14 DEGs in “Acid” set and 58 DEGs in both the “Acid” and the “Plant” set. Conversely, 57 genes exhibited differential expression in C58 but not 1D1108. Some of the notable DEGs in four major categories include: (1) up-regulated in only 1D1108: the T6SS (*impEHKLM* and *hcp*); (2) down-regulated in only 1D1108: glycine betaine/proline transport system (*proVWX*), Holliday junction DNA helicase and resolvase (*ruvABC*); (3) up-regulated in only C58: succinoglycan biosynthesis and export (*exoLTUW*); (4) down-regulated in only C58: cytochrome c oxidase (*fixOPQ*).

**FIG. 7.**
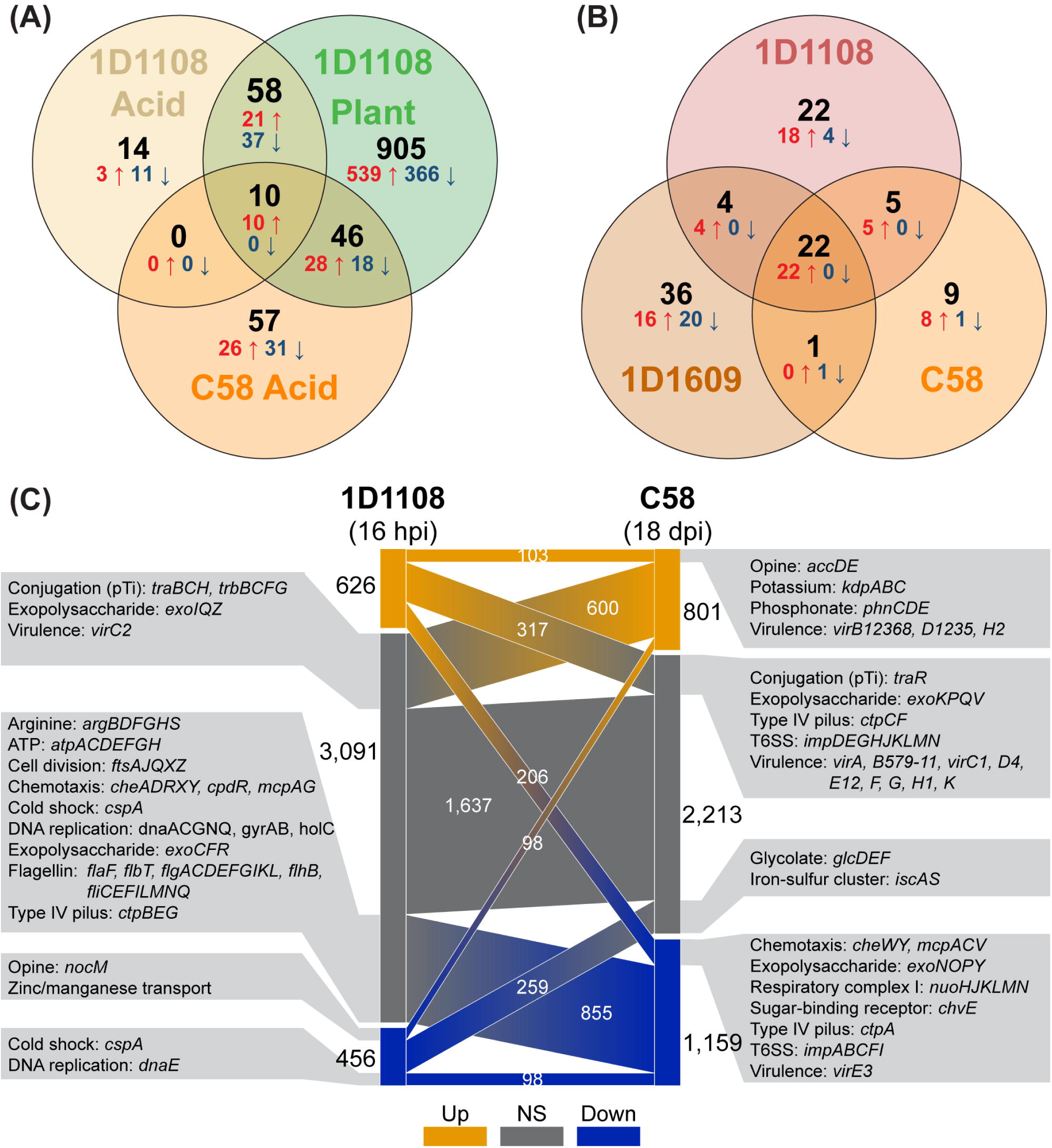
Comparisons among *Agrobacterium* transcriptome datasets. (A) Comparison to the acid treatment of strain C58 (Yuan et al., 2008). A total of 4,173 single-copy protein-coding genes were conserved between the two strains analyzed. The combined numbers of differentially expressed genes (DEGs) were indicated in black, while the numbers of up- and down-regulated genes were indicated in red and blue, respectively. (B) Comparison to the acetosyringone (AS) treatment of strains C58 and 1D1609 (Haryono et al., 2019). A total of 3,761 single-copy protein-coding genes were conserved among the three strains analyzed. (C) Comparison to the in-tumor transcriptome of C58 (González-Mula et al., 2018). The sampling time points were 16 hours post infiltration (hpi) and 18 days post infection (dpi) for strains 1D1108 and C58, respectively. The 4,173 single-copy protein-coding genes conserved between these two strains were separated into up-regulated (orange), no significant change (gray), and down-regulated (blue) in the two datasets. For each category, the gene count and notable genes were labeled.

The second comparison examined the response to the phenolic compound AS, a known inducer of *vir* gene expression. The reference study is our previous work that used the same experimental design to investigate AS-induced expression in strains C58 and 1D1609 (Haryono et al., 2019). By including 1D1108 in the comparison, the three strains share 3,761 single-copy protein-coding genes. Only 22 DEGs were consistently up-regulated (**Fig. 7B**), 20 of which belonged to the *vir* regulon (**Table S3**). The remaining 77 DEGs showed strain-specific regulation: 67 were uniquely regulated in a single strain, and 10 shared the same direction of regulation in two strains but not all three. These genes varied widely in function and genomic location (**Table S3**). Together, these two cross-strain comparisons under axenic conditions highlight substantial divergence in gene regulation, even in response to shared abiotic cues.

In the third comparison, we aimed to investigate the transcriptional regulation inside host plants. However, such datasets were rare; only one suitable study was found, which investigated C58 inside *Arabidopsis thaliana* tumors at 18 days post infection (dpi) based on microarrays (González-Mula et al., 2018). As observed in the two comparisons of axenic conditions, DEGs sharing the same patterns of regulation were rare (**Fig. 7C**). Among the 4,173 conserved homologs, only 201 showed consistent regulation, while 304 were regulated in opposite directions. Some differences may reflect species- or strain-specific regulation, but others likely result from differences in experimental design, including host plant species, tissue type, and time point. Nevertheless, some general observations can be made. For example, several genes corresponding to the *vir* regulon (*virB12368*, *virD1235*, and *virH2*) and opine transporter (*accDE*) were consistently up-regulated (**Table S3**), in line with the understanding of *Agrobacterium* ecology that these bacteria transform their hosts to obtain opines as a nutrient source (Nester, 2015). Additionally, genes involved in the transport of potassium and phosphonate were consistently up-regulated, suggesting that these two substrates may be important for agrobacterial physiology inside plants.

The T6SS showed a contrasting pattern between datasets. Most T6SS genes were up-regulated in the 1D1108 in-leaf dataset, but were either unchanged (*impDEGHJKLMN*) or down-regulated (*impABCFI*) in the C58 in-tumor dataset. These differences may reflect the stage of infection. At 16 hours post infiltration (hpi), the T6SS may be deployed to compete with other bacteria. In contrast, at 18 dpi, agrobacteria may already dominate the tumor microbiome, at least in small spatial scales, and repression of the T6SS activation could reduce costs. This interpretation is consistent with previous findings that the T6SS confers competitive advantages early during infection inside agroinfiltrated leave of *N. benthamiana* within 24 hpi (Ma et al., 2014; Wu et al., 2019), but does not play a major role in shaping tumor microbiome at 60 dpi (Wang et al., 2023).

Finally, several genes associated with chemotaxis, exopolysaccharide production, and type IV pilus biogenesis were up-regulated only in 1D1108 during early plant colonization, but not in the C58 in-tumor dataset. These genes may contribute to processes important at early stages of infection, such as surface sensing, attachment, or migration within host tissue.

### Limited in planta transcriptional overlap with *Pseudomonas syringae*

To assess whether plant-associated bacteria exhibit conserved transcriptomic responses during host colonization, we extended our comparative analysis to include two RNA-Seq datasets of *Pseudomonas syringae* pv. *tomato* strain DC3000 (Lovelace et al., 2018; Nobori et al., 2018). Although *Agrobacterium* and *Pseudomonas* differ substantially in taxonomy, ecology, and infection strategy, both colonize plant apoplast and interact with host-derived cues during the early stages of infection. Therefore, a cross-species comparison provides an opportunity to assess whether common transcriptional responses are activated by plant-derived environments. Although evolutionary divergence and methodological variation pose challenges, identifying shared expression patterns could highlight convergent adaptations or core functional themes in host colonization.

For this analysis, we identified 1,441 single-copy homologs shared between 1D1108 and DC3000, accounting for approximately one-quarter of the protein-coding genes in each genome (**Fig. 8; Table S3**). The two DC3000 datasets included for comparison were based on infiltration into *Arabidopsis thaliana* leaves, with RNA samples collected at 6 and 5 hpi, respectively. Thus, both the host plant species and the sampling time points differ from those used in the 1D1108 dataset presented in this study. Nevertheless, the same criteria of DEG inference were used.

**FIG. 8.**
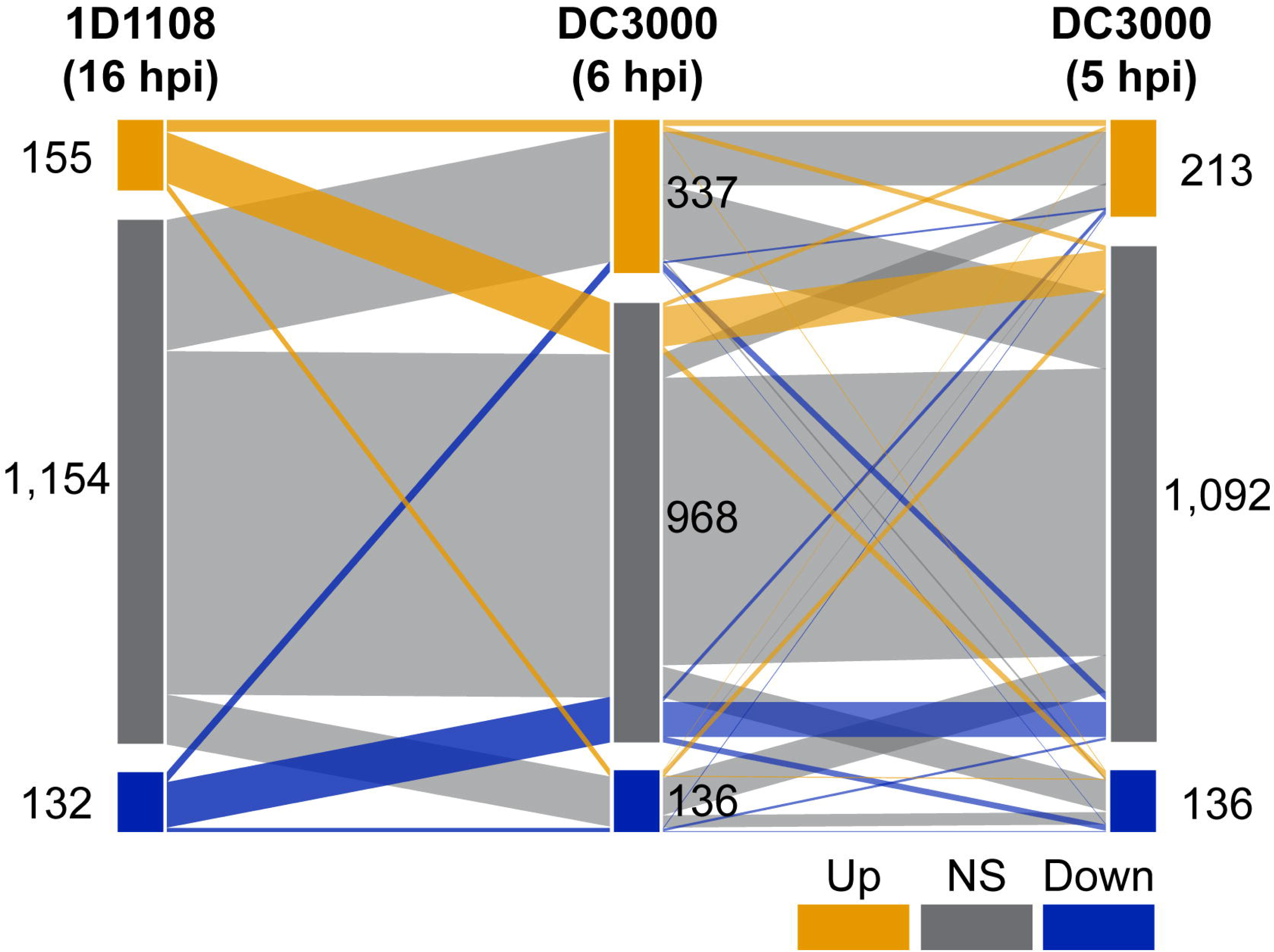
Comparisons with *planta* transcriptomes of *Pseudomonas* strain DC3000. The *in planta* transcriptome datasets of *Pseudomonas syringae* pv. tomato DC3000 in *Arabidopsis thaliana* leaves at 6 hpi (hpi) (Nobori et al., 2018) and 5 hours post infiltration (Lovelace et al., 2018) were included for comparison. The 1,441 protein-coding genes conserved between *Agrobacterium* 1D1108 and *Pseudomonas* DC3000 were separated into up-regulated (orange), no significant expression change (gray), and down-regulated (blue). The numbers in each category and correspondence between datasets were labeled.

The analysis revealed very limited congruence among the three *in planta* transcriptomes. Even between the two DC3000 datasets, which differed only slightly in the sampling time, only 135 genes were consistently up-regulated and 31 down-regulated (**Table S3**). When 1D1108 was included, the number of shared up-regulated DEGs dropped to 13, and only two genes were consistently down-regulated (**Table S3**). Despite the limited overlap, the shared up-regulated genes point to a coherent response involving central metabolism, nutrient uptake, and stress adaptation. These included genes involved in central carbon metabolism (*accB*, *ilvI*, and *sucB*), sugar utilization (*cscA* and *thuE*), amino acid and sulfur assimilation (*argJ* and *cysP*), osmoprotectant transporters (*opuBD* and *proW*), and the oxidative stress response (*ohr*). This pattern suggests that the early colonization involves metabolic reprogramming, nutrient scavenging, and defenses against host-derived stresses. In comparison, the two commonly down-regulated DEGs, acyl-CoA dehydrogenase (*acd*) and the accessory factor of xanthine dehydrogenase (*xdhC*), are both associated with energy metabolism. This finding suggests a suppression of purine catabolism and fatty acid degradation pathways during early infection, possibly reflecting a shift toward host-derived carbohydrates and osmolytes as primary energy sources.

Despite these shared elements, the overall low overlap, particularly between the two *Pseudomonas* datasets with similar designs, highlights substantial transcriptomic divergence across studies. Differences in host species, localized microenvironments, and even slight shifts in sampling time can all contribute to these discrepancies. These results underscore the difficulty of identifying a universal *in planta* transcriptional program across diverse study systems of bacterial strains and their hosts.

In contrast to the variability observed across studies, our own transcriptomic datasets showed high internal consistency and clear treatment-specific signatures (**Fig. 3**). The broader divergence among datasets likely reflects fundamental biological differences. *Agrobacterium* is a biotrophic pathogen that delivers T-DNA and effectors through its T4SS, whereas *P. syringae* employs a type III secretion system (T3SS) and associated effectors to suppress host immunity during hemibiotrophic infection. Despite these mechanistic differences, both systems were transcriptionally up-regulated *in planta* (**Table S3**), emphasizing active secretion as a shared feature of early host colonization. These findings suggest that while the transcriptional programs deployed by different plant-associated bacteria are highly context-dependent, they converge on common functional imperatives such as resource acquisition, competition, and host interface modulation.

## CONCLUSIONS

This study integrates comparative phenotyping and transcriptomic profiling to examine how environmental and host-associated cues shape gene expression in *Agrobacterium*. We focused on the wild-type strain 1D1108, which exhibited high transformation efficiencies across multiple assays. By combining *in vitro* treatments that simulate plant signals with *in planta* transcriptome analysis, we captured broad regulatory responses that influence *Agrobacterium*-mediated transformation and early-stage host colonization.

Our results demonstrate that the plant environment triggers widespread and coordinated gene expression changes in *Agrobacterium*, including the activation of the *vir* regulon, secretion systems, attachment mechanisms, and nutrient uptake pathways. While acidic pH and phenolic signals are established inducers of virulence, *in vitro* experiments captured only a subset of the transcriptional dynamics observed *in planta*. These findings reveal that host-derived cues elicit a more complex and integrated response, involving metabolic reprogramming and stress adaptation that support bacterial survival and interaction with host tissue.

Cross-species comparisons at within-genus and between-class levels revealed limited overlap in transcriptional responses, underscoring the context-specific and strain-dependent nature of bacterial gene regulation. These results highlight the need to study *Agrobacterium* biology in ecologically and physiologically relevant contexts, beyond simplified axenic conditions and a few model strains. Future work should incorporate time-resolved *in planta* transcriptomics, coupled with functional and multi-omics approaches, to unravel the molecular basis of strain variation, ecological adaptation, and transformation efficiency.

## MATERIALS AND METHODS

### Tumor assay for stable transformation efficiencies

The bacterial strains used are listed in **Table 1** and the procedure was based on that described previously (Hwang et al., 2013). The strains were grown at 28 °C in 2 mL of 523 medium overnight, then diluted 10-fold and subculture in fresh medium for 4 hours. The cells were collected by centrifugation at 8,000 x g for 10 minutes, then resuspended to OD_600_ 1.0 with 0.9% NaCl solution. The host plants, three-week-old kidney bean and soybean Tainan No. 7, were wounded on the stem by a sterilized needle before applying 5 µL bacterial suspension. For mock inoculation, 5 µL of autoclaved 0.9% NaCl solution was used. Results were collected at five weeks after inoculation. The tumor formation rates were calculated based on 11 to 16 plants in three batches. For tumor weight, data points that deviate from the average by more than three standard deviations were removed as outliers. The statistical significance was evaluated using one-way ANOVA; multiple comparisons were corrected with Tukey statistical hypothesis testing.

### Agroinfiltration for transient transformation efficiencies

The procedure was modified from an established protocol (Sparkes et al., 2006). The RUBY reporter system (He et al., 2020), in which the transgene expression leads to a visible pigment betalain, was chosen for quantifying the transformation efficiency. Agrobacterial strains carrying the 35S:RUBY plasmid (Addgene number 160908) were grown in 3 mL of 523 medium with spectinomycin (250 µg/mL) at 28 °C overnight, centrifuged at 10,000 x g for 10 minutes, then resuspended in infiltration buffer (10 mM MES, 10 mM MgCl_2_, 150 µM AS) with OD_600_ adjusted to 0.2. Leaves of 30-day old *N. benthamiana* were used for infiltration. For quantification, leaf discs were collected two days after infiltration and soaked in 200 µl betalain extraction buffer (10% EtOH, 0.1% formic acid). After incubation at room temperature overnight, 150 µl of the extraction buffer was used for absorbance measurement. The raw betalain accumulation level was calculated as ((OD_475_ - OD_600_) + (OD_535_ - OD_600_)). To calculate relative betalain accumulation levels, the individual measurements were normalized to values derived from the reference strain C58 obtained from the same leaf. Three batches with 8 to 10 leaves per batch were used for one-way ANOVA and Tukey post-hoc test.

### RNA-Seq experiment

The experimental design of four conditions with three biological replicates each is illustrated in **Figure 3A**. The *in vitro* experiments were conducted based on the procedure described previously (Haryono et al., 2019). Briefly, the strains were grown on 523 agar plates at 28 °C for three days, then individual colonies were picked and grown in 5 mL 523 liquid medium at 28 °C overnight. The bacterial cells were collected by centrifugation at 6,000 x g for 4 minutes, then resuspended in AB-MES medium (pH 7.0) to OD_600_ 10. For each sample, 0.1 mL of the suspension was added to 4.9 mL AB-MES (pH 7.0 or pH 5.5) and cultured at 28 °C for 6 hours before adding 5 µl of 200 µM AS dissolved in DMSO for induction or 5 µl of DMSO as control. For *in planta* experiments, bacterial cells in 15 mL of suspension in AB-MES (pH 7.0) were collected by centrifugation and resuspended in 15 mL infiltration buffer (10 mM MES, 10 mM MgCl_2_) before being infiltrated into 31-day-old *N. benthamiana*. To enrich the bacterial cells, we followed the procedure described previously (Nobori et al., 2018). The infiltrated leaves were grounded with liquid nitrogen in a mortar and incubated in 30 mL ice-cold bacterial isolation buffer (25 mM TCEP, 9.5% ethanol, 0.5% phenol, pH 4.5) at 4 °C with 100 rpm shaking for 20 hours. The buffer was filtered using 6 µm sterilized filter mesh, centrifuged at 3,200 x g and 4 °C for 20 min before removing the supernatant. The pellet was resuspended with 900 µL ice-cold bacterial isolation buffer and centrifuged at 3,800 x g and 4 °C for 20 min. After centrifugation, the upper white layer that contains enriched bacterial cells was carefully collected by pipetting into a clean 1.5 mL tube.

The RNA samples were prepared using HiYield Total RNA Extraction Kit (Cat. No. YRB50, Arrowtech, Taiwan). For each sample, the expression level of *virB1* was checked by RT-qPCR with *recA* as the internal control. The primers for these two genes are: *virB1* (forward: ACCGCGCGTCCAAAAGATGAT; reverse: ACATCCCATGTTTCCTCGGAT) and *recA* (forward: AGAGAACCCGTCGAAATGGTCT; reverse: TCGGTTCCAATGAAAACGTGGTTGA). The strand-specific RNA-Seq library preparation was processed by the core facility of Academia Sinica (Taipei, Taiwan) using the TruSeq Stranded mRNA Library Prep Kit (Cat. No. 20020594, Illumina, USA). To enrich the bacterial mRNA molecules, three additional steps were incorporated. First, for the *in planta* samples, plant mRNA molecules were removed using the RNA Purification Beads from the TruSeq Stranded mRNA Library Prep Kit, which utilize poly-T oligo-attached magnetic beads to capture the poly-A tail of eukaryotic mRNA molecules. Second, for all samples, bacterial rRNA molecules were depleted using the QIAseq FastSelect -5S/16S/23S Kit (Cat. No. 335925, QIAGEN, Aarhus, Denmark). Third, for the *in planta* samples, the depletion of plant rRNA molecules was performed using the QIAseq FastSelect –rRNA Plant Kit (Cat. No. 334315, QIAGEN, Aarhus, Denmark). After these additional processing steps, all samples underwent cDNA synthesis, adapter ligation, and library amplification using TruSeq Stranded mRNA Library Prep with SuperScript III Reverse Transcriptase (cat. No. 18080044, Thermo Fisher Scientific, United States), KAPA HiFi HotStart ReadyMix (cat. No. 07958935001/KK2602, Roche, Basel, Switzerland), and Agencourt AMPure XP (cat. No. A63881, Beckman-Coulter, Brea, CA).

For quality control, we performed DNA quantification using Qubit dsDNA HS quantification (Cat. No. Q32854, Invitrogen, United States), Fragment Analyzer DNA size profiling using HS NGS Fragment Kit (Cat. No. DNF-474-500, Agilent Technologies, United States), and digital PCR quantification using QX200 Droplet Digital PCR EvaGreen Supermix System (Cat. No. 1864001, 1864034, Bio-Rad, United States). The Illumina paired-end sequencing based on the NovaSeq 6000 platform was outsourced to Genomics BioSci and Tech Co, Ltd (New Taipei, Taiwan).

### Transcriptomics analysis

Unless stated otherwise, the bioinformatics tools were used with the default settings. The Illumina raw reads were processed using Trimmomatic v0.39 (Bolger et al., 2014) for adaptor removal and quality trimming with the settings “ILLUMINACLIP:2:30:10:1:True LEADING:20 TRAILING:20 MINLEN:50”. The processed reads were mapped to *Agrobacterium* strain 1D1108 genome (GenBank accession: GCA_003666425.1) using BWA v0.7.17 (Li and Durbin, 2009). The mapping results were processed using SAMtools v1.9 (Li et al., 2009) to calculate reads in regions corresponding to rRNA, tRNA, and protein-coding genes. For the *in planta* datasets, reads that did not map to the bacterial genome were mapped to the *N. benthamiana* genome (GCA_000723945.1) to confirm that those reads were originated from the plant hosts, rather than contaminations.

To infer the gene expression levels, the mapped reads were counted by using HTSeq v2.0.2 (Putri et al., 2022) with the settings “-t CDS -s no --nonunique all”. The result was further processed using DESeq2 v1.36.0 (Love et al., 2014) to infer the normalized expression levels based on the median-of-ratios method. The DEGs were defined based on the criteria of |log_2_(fold change)| ≥ 1 and *p*-adjusted < 0.01. To avoid spurious results, genes with total read counts < 10 were excluded. Principal component analysis was performed using the “plotPCA” function of DESeq2.

### Comparisons among transcriptomics studies

For comparisons with other transcriptomics studies, the genome sequences of *Agrobacterium* C58 (GCA_000092025.1), *Agrobacterium* 1D1609 (GCA_002943835.1), and *Pseudomonas syringae* pathovar *tomato* DC3000 (GCA_000007805.1), were obtained from GenBank. The homologous genes among 1D1108 and other bacteria were identified using OrthoMCL v1.3 (Li et al., 2003).

Based on the homologous genes, the lists of DEGs identified in this study were compared with those identified in previous works, including those that examined the effects of an acidic environment on C58 (Yuan et al., 2008), AS treatment on strains C58 and 1D1609 (Haryono et al., 2019), the in-tumor environment on C58 (González-Mula et al., 2018), and the in-leaf environment on DC3000 (Lovelace et al., 2018; Nobori et al., 2018). To maintain consistency, the lists of DEGs were all filtered based on the same criteria of |log_2_(fold change)| ≥ 1 and *p*-adjusted (or equivalent when available) < 0.01. Because the AS treatment study on C58 and 1D1609 (Haryono et al., 2019) was conducted by our team, we re-analyzed the raw reads using the same bioinformatic procedure as described in this study. For other comparisons, we obtained the lists of DEGs from the supplementary materials of previous publications (Yuan et al., 2008; González-Mula et al., 2018; Lovelace et al., 2018; Nobori et al., 2018).

## DATA AVAILABILITY

All Illumina RNA-Seq datasets are available in the National Center for Biotechnology Information (NCBI) under BioProject accession PRJNA1111437.

## AUTHOR CONTRIBUTION

Conceptualization: YW, EML, CHK.

Funding acquisition: CHW, EML, CHK.

Investigation: YW, HYC.

Methodology: YW, HYC, CHW, EML, CHK.

Project administration: EML, CHK.

Supervision: EML, CHK.

Validation: YW, HYC, EML, CHK.

Visualization: YW, HYC, CHK.

Writing – original draft: YW, CHK.

Writing – review & editing: YW, CHW, EML, CHK.

## USAGE OF ARTIFICIAL INTELLIGENCE TOOLS

ChatGPT-4o was used to assist with brainstorming manuscript organization, suggesting improvements in wording, and correcting grammar. The vast majority of the writing and the final text are the authors’ original work. No generative AI tools were used in the preparation of figures or tables.

## ACKNOWLEDGMENTS

We thank members of the Agrobacteria-Mediated Transformation Team and the Plant and Environmental Microbiology Group in our institute for scientific discussions. Technical assistance was provided by Shu-Jen Chou, Ai-Ping Chen, Mei-Jane Fang, Ming-Ling Cheng, and Ching-I Kuo. The Illumina sequencing library preparation was carried out by the Genomic Technology Core (Institute of Plant and Microbial Biology, Academia Sinica). The Illumina sequencing service was provided by Genomics BioSci and Tech Co, Ltd (New Taipei, Taiwan).

## SUPPLEMENTAL MATERIAL

**Table S1**: Summary statistics of the RNA-Seq experiment.

**Table S2**: Detailed results of the RNA-Seq experiment. For each gene, the normalized expression levels in all 12 samples and the patterns of differential expression in the three pairwise comparisons were inferred based on DESeq2.

**Table S3**: Lists of transcriptomic comparisons. (A) Comparison to the acid treatment of strain C58 (Yuan et al., 2008). (B) Comparison to the acetosyringone (AS) treatment of strains C58 and 1D1609 (Haryono et al., 2019). (C) Comparison to the in-tumor transcriptome of C58 (González-Mula et al., 2018). (D) Comparison to the *in planta* transcriptome datasets of *Pseudomonas syringae* pv. tomato DC3000 in *Arabidopsis thaliana* leaves at 5 hours post infiltration (hpi) (Lovelace et al., 2018) and 6 hpi (Nobori et al., 2018).

